# Conformational plasticity of the human norovirus GII.3 capsid reveals alternative P domain interaction networks

**DOI:** 10.64898/2026.06.25.734440

**Authors:** Chihong Song, Motohiro Miki, Reiko Takai-Todaka, Kosuke Murakami, Kazuhiko Katayama, Kazuyoshi Murata

**Author notes:** Corresponding authors: (KM), (KK).

## Abstract

Human noroviruses (HuNoVs) are a leading cause of acute gastroenteritis worldwide, yet no effective antiviral therapeutics are currently available. Although environmentally induced capsid conformational changes associated with infectivity have been reported in murine noroviruses (MNVs), comparable conformational switching has not been demonstrated in HuNoVs. In this study, we generated HuNoV GII.3 virus-like particles (VLPs) using a baculovirus expression system and identified two distinct T = 3 particle populations coexisting within VLP preparations derived from a single strain through cryo-electron microscopy single-particle analysis. Comparative structural analysis revealed that these two T = 3 capsid conformations correspond to the resting and rising states of the protruding (P) domain. Rearrangement of the P domain alters intermolecular interactions between adjacent capsid subunits, resulting in distinct capsid surface architectures. In the resting state, intermolecular contacts were mediated predominantly by the P2 subdomain, with limited contribution from the P1 subdomain. In contrast, the rising state exhibited a shift toward an alternative interaction interface primarily involving the P1 subdomain. These findings demonstrate previously unrecognized structural polymorphism in the HuNoV capsid and provide evidence that conformational switching may occur in HuNoVs. Our results offer new insights into norovirus capsid dynamics and may inform future structure-based vaccine and antiviral drug development.

## 1. Introduction

Human norovirus (HuNoV) is a leading cause of nonbacterial acute gastroenteritis worldwide and is responsible for frequent outbreaks in crowded or semi-enclosed environments, including long-term care facilities, hospitals, and childcare centers [1–3]. Globally, HuNoV is estimated to cause approximately 600-700 million infections and nearly 200,000 deaths each year, representing a substantial public health burden [1,4].

Noroviruses (NoVs) are non-enveloped, positive-sense single-stranded RNA viruses belonging to the family *Caliciviridae* [5]. Their ∼7.5 kb genome contains three open reading frames (ORF1-3). ORF1 encodes the non-structural proteins required for viral replication, ORF2 encodes the major capsid protein VP1, and ORF3 encodes the minor structural protein VP2, which is located within the capsid. Both VP1 and VP2 are translated from a subgenomic RNA [5]. Because efficient propagation of HuNoV in conventional cell culture system remains challenging and infectious virus handling poses biosafety concerns, structural and immunological studies have relied largely on virus-like particles (VLPs) assembled from recombinant VP1. These genome-free particles provide a safe and versatile platform for structural characterization of the viral capsid and have also served as important tools for antigen presentation and vaccine development [1,4].

The canonical calicivirus capsid exhibits a T = 3 icosahedral architecture composed of 180 copies of VP1, first resolved at high resolution through X-ray crystallography and cryo-electron microscopy (cryo-EM) studies of Norwalk virus (GI.1) VLPs [6]. Within the T = 3 lattice, VP1 occupies three quasi-equivalent conformations, designated subunits A, B, and C. A/B dimers are located around the 5-fold symmetry axes, whereas C/C dimers are positioned on the 2-fold axes. Structurally, VP1 consists of a flexible N-terminal domain (NTD), a conserved shell (S) domain that forms the capsid shell, and an outward-facing protruding (P) domain connected to the S domain through a flexible hinge region. The P domain is further divided into the lower P1 and upper P2 subdomains, with the P2 subdomain playing a central role in receptor interactions and antigenic variation [5,6]. Consistent with this role, antigenic drift and immune escape predominantly occur on the P2 surface, whereas the S domain remains relatively conserved across members of the *Caliciviridae* family [4].

Recent studies have revealed that norovirus capsids are dynamic structures capable of undergoing environmentally regulated conformational transitions. In murine norovirus (MNV), changes in pH and metal-ion concentration induce a reversible ∼70° rotation of the P domain, generating either a resting state in which the P domain lies close to the shell surface or a rising state in which it is elevated away from the shell [7]. The resting state has been associated with enhanced receptor engagement and infectivity, whereas the biological significance of the rising state remains less well understood and may relate to environmentally responsive regulation of infectivity or immune evasion [7–9]. Furthermore, Sherman et al. demonstrated that low pH, bile salts, and divalent metal ions promote contraction of the P domain toward the shell, and that metal-ion-induced activation can be reversed by EDTA treatment [9].

Capsid conformational plasticity has also emerged as an important factor in HuNoV biology and immune recognition [10]. The broadly reactive llama-derived nanobody M4 neutralizes multiple GII.4 variants by targeting a conserved epitope that becomes fully accessible in a raised VP1 conformation and can additionally promote capsid destabilization and particle disassembly [11]. Interestingly, although M4 binds to the P domains of both GII.4 and GII.3 viruses and induces disassembly of GII.3 VLPs, neutralization was observed only for GII.4 viruses in human intestinal enteroid infection assays [11]. These findings suggest genotype-specific differences in the capsid stability and conformational dynamics and highlight the importance of characterizing the structural properties of GII.3 viruses. GII.3 strains are frequently associated with pediatric gastroenteritis and sporadic outbreaks, and recent epidemiological evidence indicates their increasing prevalence in certain regions. Notably, GII.3[P12] was identified as the predominant outbreak-associated genotype in Beijing, China, during 2021-2023, where genomic analyses revealed ongoing amino-acid substitution-driven evolution and regional transmission dynamics [12].

Despite growing evidence for capsid plasticity in both murine and human noroviruses, direct structural evidence for conformational switching within HuNoV particles remains limited. In the present study, we produced HuNoV GII.3 VLPs using a baculovirus expression system and characterized their structures by cryo-EM single-particle analysis. We identified two distinct T = 3 capsid conformations corresponding to the resting and rising states of the P domain. Comparative structural analysis revealed that transitions between these states reorganize intermolecular interactions between neighboring capsid subunits, generating alternative capsid surface architectures. These findings provide direct structural evidence for conformational polymorphism in HuNoV capsids and offer insights into the molecular basis of capsid plasticity, with potential implications for vaccine design and antiviral development.

## 2. Results

### 2.1 Single particle analysis and structural characterization of HuNoV GII.3 VLPs

HuNoV GII.3 VLPs were produced using a baculovirus expression system. Cryo-EM micrographs of the purified sample revealed heterogeneous particle populations of varying sizes, which were presumed to correspond to T = 1, T = 3, and T = 4 capsids (Fig. S1). Among these, T = 3 particles with a diameter of approximately 40 nm were selected by automated particle picking and subjected to single-particle analysis. 2D classification revealed two distinct particle populations that differed in the position of the P domain (Fig. 1A, B and Fig. S1). The first population exhibited a conformation in which the P domain was positioned close to the S-domain shell. This structure was reconstructed at a resolution of 2.8 Å. The S-domain shell showed high local resolution (approximately 2.7-2.9 Å), whereas the P domain region was resolved at relatively lower resolution (Fig. 1C and Fig. S1). The second population exhibited a conformation in which the P domain was elevated and detached from the shell. This structure was reconstructed at 7.2 Å resolution, with the P domain region displaying particularly low local resolution (∼10 Å), indicative of increased structural flexibility (Fig. 1D and Fig. S1). These two conformations correspond to the resting and rising states previously reported for murine norovirus [7].

**Figure 1.**
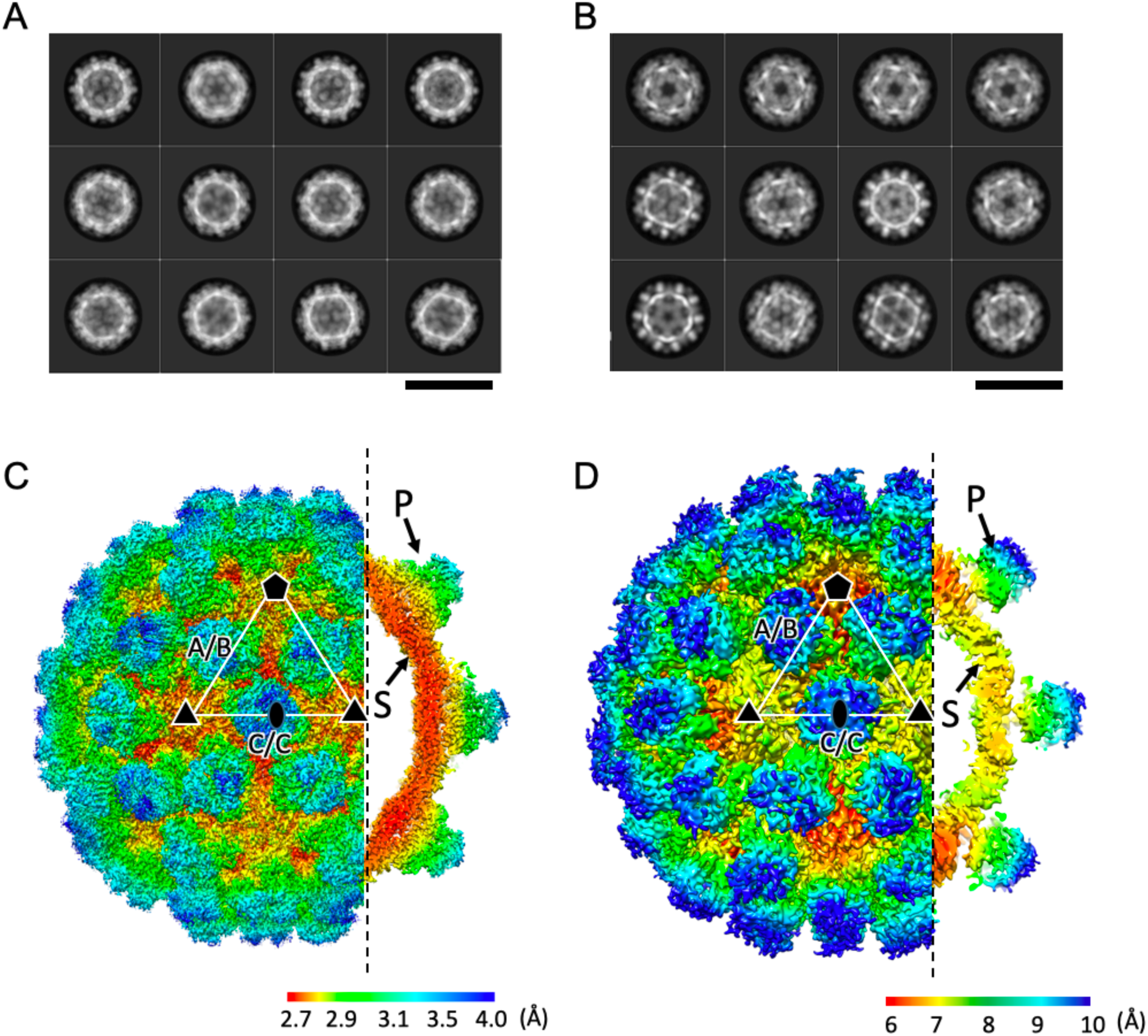
Cryo-EM structures of HuNoV GII.3 VLPs. (A) Representative 2D class averages of HuNoV GII.3 T = 3 VLPs exhibiting the resting P domain conformation. (B) Representative 2D class averages of HuNoV GII.3 T = 3 VLPs exhibiting the rising P domain conformation. (C) Cryo-EM map of the HuNoV GII.3 T = 3 VLP in the resting state, colored according to local resolution. The cutaway view reveals that the P domains are positioned close to the S-domain shell. (D) Cryo-EM map of the HuNoV GII.3 T = 3 VLP in the rising state, colored according to local resolution. The cutaway view shows that the P domains are elevated above the S-domain shell. The locations of the A/B and C/C dimers within a single icosahedral asymmetric unit are indicated. Scale bars in (A) and (B) are 40 nm. The color codes of local resolution are indicated in C and D, respectively.

Because the P domain was less well resolved than the shell in the full-particle reconstructions, focused classification and refinement were performed following symmetry expansion to better characterize the structure and dynamics of the P domain dimers. Symmetry-expanded particles were subjected to focused 3D classification without alignment, allowing each class to retain the spatial position of the corresponding P domain dimer within the T = 3 capsid.

For the resting state, focused refinement improved the resolutions of the C/C and A/B dimeric P domains to 3.1 Å and 3.4 Å, respectively (Fig. S2). 3D classification revealed that the C/C dimer exhibited positional variation of up to approximately 9 Å toward the region lacking an adjacent A/B dimer, whereas displacement of the A/B dimer was limited to approximately 3 Å. These findings indicate that the degree of P domain flexibility differs among quasi-equivalent positions even within the resting capsid (Fig. S2). Focused refinement of the P domain in the rising state yielded a reconstruction at 4.7 Å resolution (Fig. S3). 3D classification revealed positional variation of approximately 11 Å among different classes. Unlike the resting state, this movement was not confined to a specific direction but occurred in multiple directions. These observations suggest that the P domain possesses greater spatial freedom in the rising state, likely as a consequence of reduced interactions with both the S-domain shell and neighboring P domain dimers.

Collectively, these results demonstrate that the T = 3 capsid of HuNoV GII.3 VLPs adopts two distinct conformational states: a resting state, in which the P domain is positioned close to the shell, and a rising state, in which the P domain is elevated away from the shell. Notably, the rising state exhibits substantially greater positional heterogeneity of the P domain, indicating enhanced structural flexibility.

### 2.2 Model building of HuNoV GII.3 capsids

The resting T = 3 HuNoV GII.3 VLP was reconstructed at 2.8 Å resolution, providing sufficient map quality for atomic model building of the VP1 capsid protein (Fig. 2A-C). The asymmetric unit of the T = 3 capsid consists of three quasi-equivalent VP1 subunits, designated A, B, and C. Each VP1 subunit comprises an S domain, which forms the continuous capsid shell, and a protruding P domain. The atomic model of the resting state fitted well into the cryo-EM density map (Fig. 2A). Most secondary-structure elements in both the S and P domains were clearly resolved, and side-chain densities were visible even within the P domain despite its relatively lower local resolution, supporting the accuracy of the model (Fig. 2B). In contrast, the density corresponding to residues 304-307 at the apex of the P2 subdomain was weak and discontinuous, preventing reliable modeling of this region.

**Figure 2.**
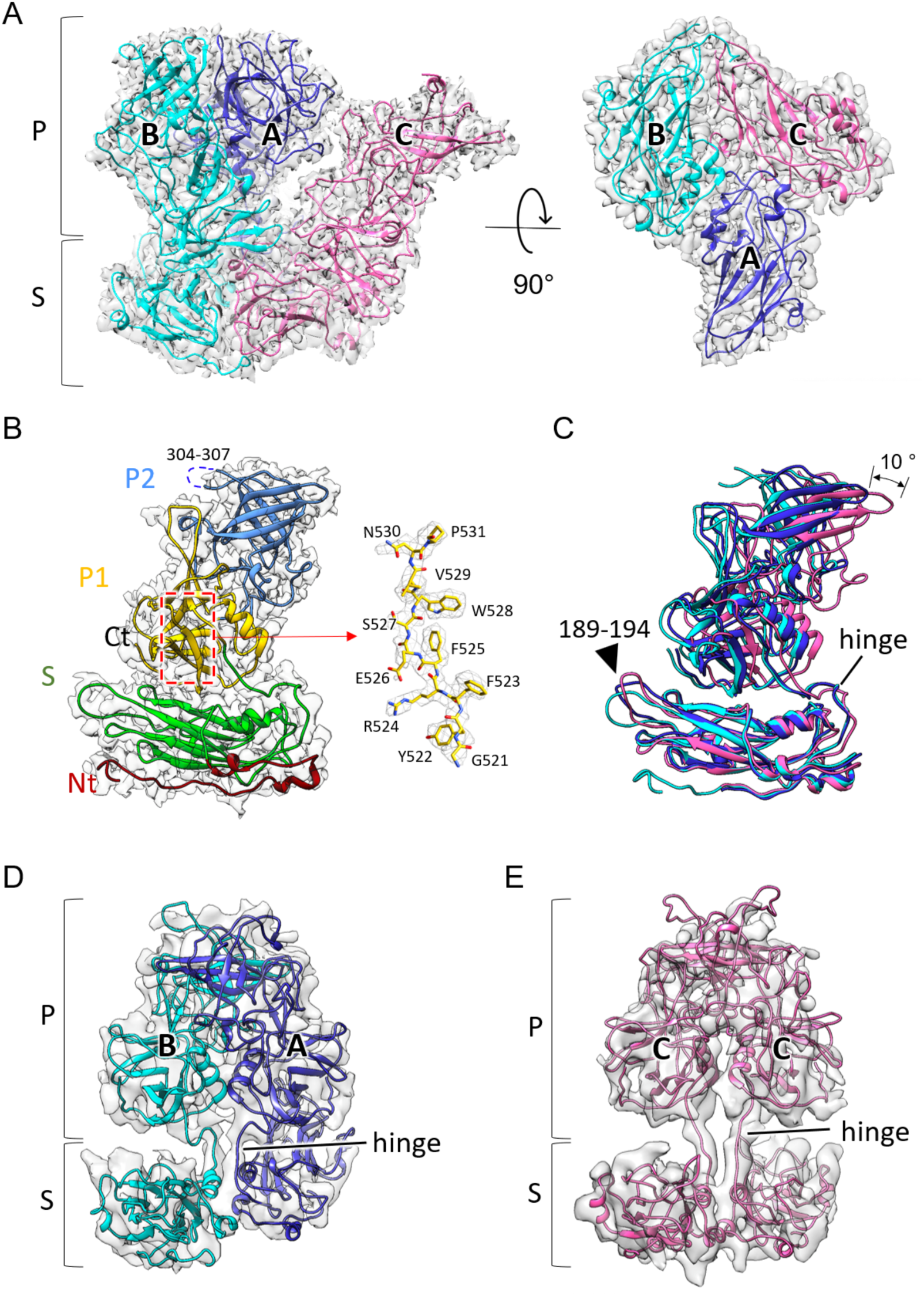
Atomic Model of the HuNoV GII.3 T = 3 capsid and structural comparison of resting and rising states. (A) Atomic model of the resting state HuNoV GII.3 T = 3 capsid fitted into the cryo-EM map. The quasi-equivalent A, B and C subunits are colored blue, cyan, and pink, respectively. (B) Structure of the VP1 capsid protein in the resting state, highlighting the N-terminal region, S domain, P1 subdomain, and P2 subdomain. The enlarged view shows a representative region of the atomic model fitted into the cryo-EM map. Map corresponding to residues 304-307 was poorly resolved, preventing reliable modeling of this loop region. (C) Superposition of the A, B, and C subunits aligned using the S domain as a reference. Minor conformational differences are observed in the region encompassing residues 189-194. (D, E) Models of the rising state A/B and C/C dimers generated by rigid-body fitting of the resting state structure into the corresponding cryo-EM maps, followed by local refinement of the hinge region.

To examine conformational differences among the three quasi-equivalent VP1 subunits, the A, B, and C subunits were superimposed using the S domain as a reference (Fig. 2C). The S domains aligned closely overall, although local structural differences were observed in the loop comprising residues 189-194. Comparison of the P domain positions following S-domain alignment revealed distinct orientations among the subunits. Notably, the position of the A/B and C subunits differed by approximately 10° in orientation, with the lower position of the hinge region appearing to serve as the pivot point for this rotation.

Because the rising state reconstruction was limited to 7.2 Å resolution, *de novo* atomic model building was not feasible. Instead, a model of the rising conformation was generated by fitting the resting state model into the corresponding density map. To accomplish this, the S and P domains of the resting state model were separated and independently fitted into the densities of the rising state reconstruction as A/B and C/C dimers (Fig. 2D, E). The hinge region connecting the S and P domains was subsequently adjusted based on the map density.

The resulting model indicates that the resting and rising states share a similar overall capsid architecture but differ substantially in the position of the P domain. These observations suggest that the transition between the two conformational states is primarily mediated by movement of the P domain around the hinge region connecting the S and P domains.

### 2.3 Rearrangement of P domain interactions in the resting and rising states

To further characterize the structural differences between the resting and rising states, we compared the positions and orientations of the P domains in the A/B and C/C dimers of HuNoV GII.3 VLPs. In the resting state, the P domains of both dimers were located close to the S-domain shell, whereas in the rising state they were displaced outward from the capsid surface (Fig. 3A, B). In the A/B dimer, the P domain underwent an approximately 55° rotation relative to the resting conformation and was elevated by ∼1.0 nm from the shell surface (Fig. 3A-B). This movement was accompanied by a slightly inward tilt of the dimer axis toward the 5-fold axis by approximately 11° (Fig. 3B). The C/C dimer similarly exhibited an approximately 55° rotation and a 1.3 nm elevation of the P domain. However, unlike the A/B dimer, the C/C dimer showed little apparent tilting, with the P domain moving almost vertically away from the shell.

**Figure 3.**
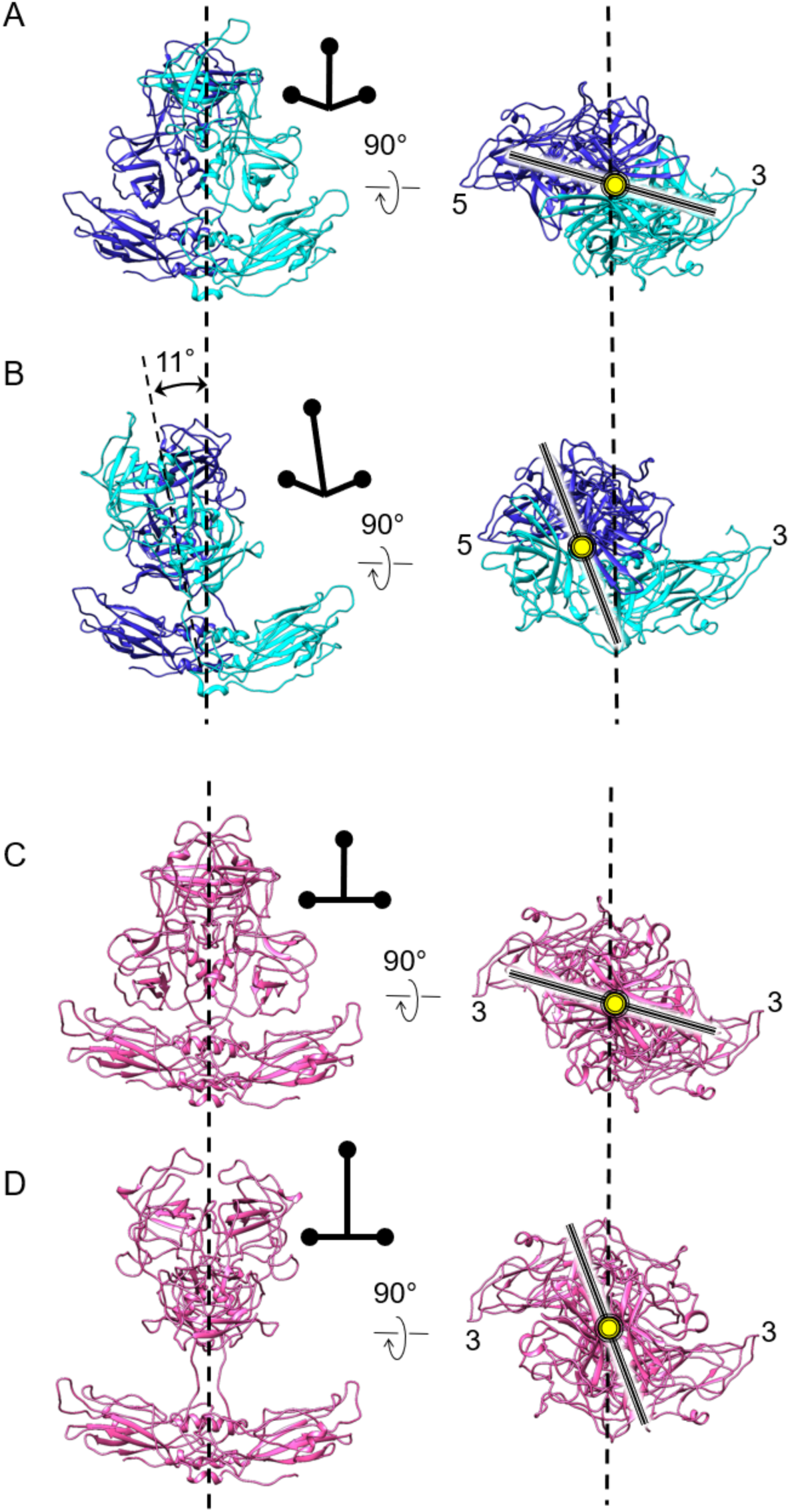
Positional changes of the P domains in the resting and rising states. (A, B) Side and top views of the A/B dimer in the resting (A) and rising (B) states. In the rising state, the P domain is elevated above the S-domain shell and undergoes an approximately 55° rotation, accompanied by an approximately 11° tilt of the dimer axis. (C, D) Side and top views of the C/C dimer in the resting (C) and rising (D) states. Similar to the A/B dimer, the C/C dimer exhibits elevation and an approximately 55° rotation of the P domain in the rising state. However, no appreciable tilt of the dimer axis is observed.

We next examined how this conformational change affects intermolecular interactions between neighboring P domains (Fig. 4). In the resting state, adjacent P domains formed extensive contacts involving both the P1 and P2 subdomains, with particularly prominent interactions occurring within the outer P2 region (Fig. 4A-D). The C/C dimer was surrounded by four A/B dimers and formed two distinct types of interaction interfaces depending on the orientation of the neighboring A/B dimers (Fig. 4A-C). At the interface formed by the A/B_1_, C/C, and A/B_4_ dimers, residues Lys375, Asp330, Met424, Asp489, and Thr490 of the A/B dimer were positioned in close proximity to Phe420, His422, Met424, Asn433, and Pro435 of the C/C dimer, forming a network of hydrophobic and electrostatic interactions (Fig. 4D). Similar interaction patterns were observed at the interface involving the A/B_2_, C/C, and A/B_3_ dimers. These P2-mediated contacts appear to tightly link neighboring P domains and contribute to maintenance of the compact resting state architecture.

**Figure 4.**
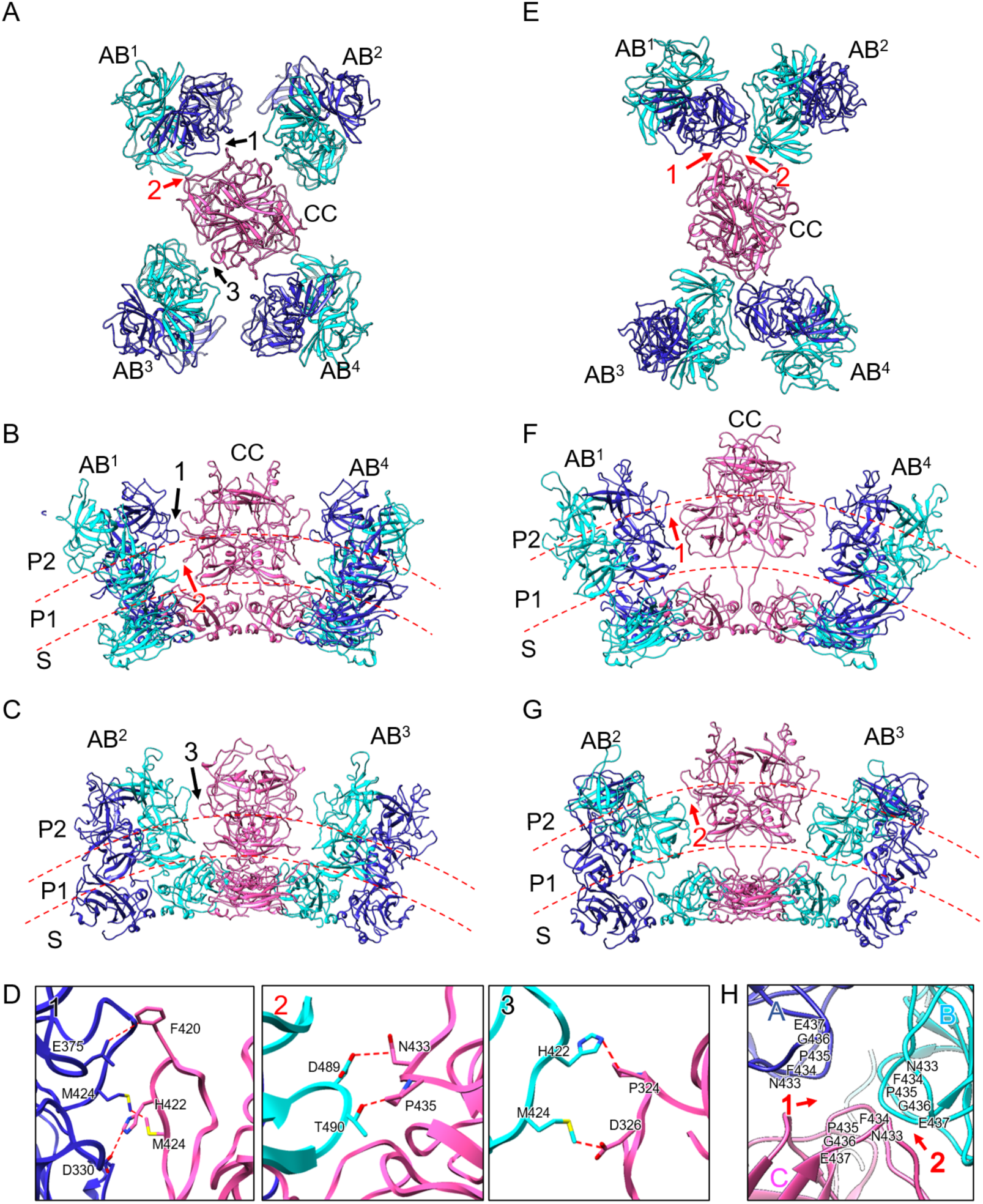
Rearrangement of intermolecular P domain interactions between the resting and rising states. (A-D) Intermolecular interactions among P domains in the resting state. (A) Top view of a central C/C dimer surrounded by four neighboring A/B dimers (AB¹-AB⁴). (B, C) Side views of the AB¹-C/C-AB⁴ and AB²-C/C-AB³ interfaces, respectively. The S domain, P1 subdomain, and P2 subdomain are indicated. (D) Enlarged views of the interfaces highlighted in (A-C), showing representative residues involved in intermolecular contacts between neighboring P domains. (E-H) Intermolecular interactions among P domains in the rising state. (E) Top view of a central C/C dimer surrounded by four neighboring A/B dimers (AB¹-AB⁴). (F, G) Side views of the AB¹-C/C-AB⁴ and AB²-C/C-AB³ interfaces, respectively. In the rising state, the P domains are elevated above the S-domain shell, and the interaction interfaces are shifted toward the P1 subdomain. (H) Enlarged view of the interfaces highlighted in (E-G), showing representative residues involved in P1-mediated intermolecular contacts between neighboring P domains.

In the rising state, the P domain interaction network was markedly reorganized. Elevation of the P domains from the S-domain shell disrupted the P2-mediated contacts observed in the resting state (Fig. 4E-H). Instead, intermolecular interactions were concentrated within the P1 subdomain, where an extended loop comprising residues 433-437 formed contacts between adjacent A/B and C/C dimers (Fig. 4E). Thus, whereas the resting state is stabilized by extensive lateral interactions mediated primarily through the P2 subdomain, the rising state is maintained by P1-mediated contacts following disruption of the P2 interface.

Together, these observations demonstrate that the transition from the resting to the rising state involves not only elevation and rotation of the P domain but also a substantial reorganization of the intermolecular interaction network. In the resting state, P2-mediated contacts constrain neighboring P domains in a compact arrangement near the capsid surface. In contrast, the interaction interface shifts toward the P1 region in the rising state, allowing the elevated P domains to remain connected while adopting a more open and flexible configuration.

## 3. Discussion

Although structural studies of the GII.3 genotype have been relatively limited compared with those of GII.4, recent reports of increasing outbreaks caused by GII.3[P12] noroviruses and whole-genome analyses have highlighted the need for a better understanding of the structural and functional properties of this genotype [12]. In the present study, cryo-EM analysis of HuNoV GII.3 VLPs revealed that the P domains of the T = 3 capsid adopt two distinct conformational states, referred to as the resting and rising states. The coexistence of these conformations within the same genotype, sample, and capsid geometry demonstrated that the HuNoV GII.3 capsid possesses intrinsic conformational plasticity. Similar to the conformational transitions reported for murine norovirus (MNV), these observations suggest that human norovirus capsids can also populate multiple metastable structural states.

In MNV, reversible capsid conformational changes are induced by environmental factors such as bile acids, metal ions, and pH, and have been linked to receptor binding, infectivity, and immune evasion [7–9,13,14]. In contrast, direct structural evidence for comparable conformational switching in HuNoV has remained limited because infectious particles are difficult to culture and purify in sufficient quantities for structural analysis. The identification of both resting and rising conformations in HuNoV GII.3 therefore supports the notion that capsid conformational plasticity is conserved across human and murine noroviruses.

A key finding of this study is that the resting and rising states differ not only in the height of the P domain, but also in the organization of intermolecular interactions between neighboring P domains. In the resting state, adjacent P domains were connected primarily through contacts involving the P2 subdomain. These interactions linked neighboring A/B and C/C dimers and appeared to constrain the P domains in a compact arrangement near the capsid surface. In contrast, the rising state lacked these P2-mediated contacts, and the interaction interface shifted toward the P1 subdomain. Thus, the transition from the resting to the rising state involves both large-scale repositioning of the P domains and extensive remodeling of the interaction network that stabilizes each conformational state.

To place these observations in a broader context, we compared the resting and rising structures of HuNoV GII.3 with those reported for MNV (Fig. 5). In the resting state, the C/C dimer in both viruses interacted with surrounding A/B dimers primarily through the P2 subdomain. However, whereas the MNV C/C dimer contacted mainly two neighboring A/B dimers, the HuNoV GII.3 C/C dimer formed a more extensive interaction network involving four surrounding A/B dimers (Fig. 5A, B). These observations suggest that P domains of HuNoV GII.3 in the resting state are more densely interconnected on the capsid surface. In the rising state, by contrast, P2-mediated interactions were disrupted in both viruses and replaced by contacts centered on the P1 subdomain (Fig. 5C, D). Thus, although HuNoV and MNV share a common mechanism involving P domain elevation and interface switching, they differ in the complexity of the resting state interaction network.

**Figure 5.**
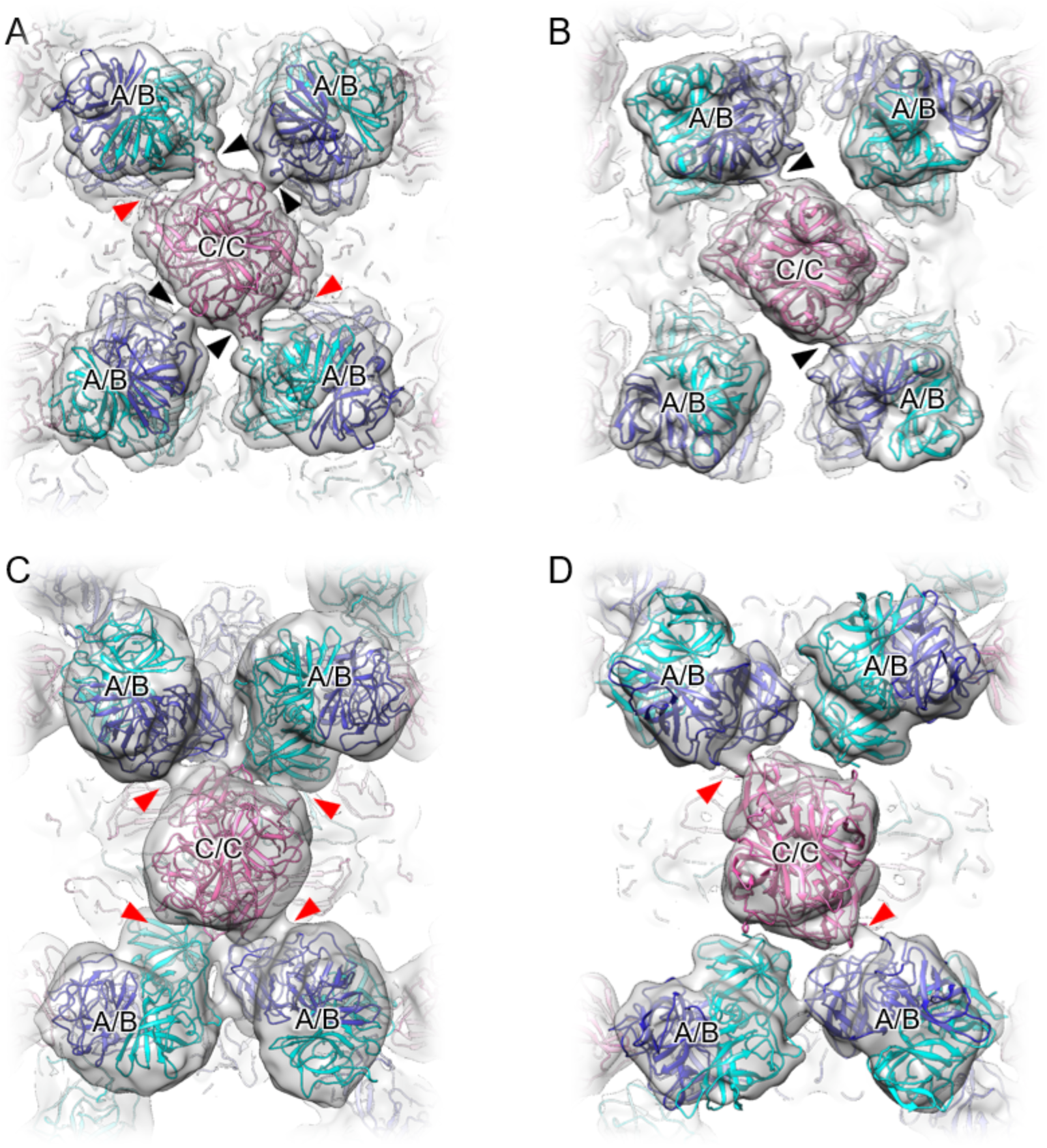
Comparison of P domain interaction networks between HuNoV and MNV. (A, B) P domain interaction networks in the resting states of HuNoV GII.3 (A) and MNV (B). (C, D) P domain interaction networks in the rising states of HuNoV GII.3 (C) and MNV (D). The central C/C dimer and the surrounding A/B dimers are shown as models fitted into the corresponding cryo-EM maps. Black arrowheads indicate P2-mediated intermolecular interactions, whereas red arrowheads indicate P1-mediated intermolecular interactions between neighboring P domains.

The lower resolution of the rising state reconstruction further supports the dynamic nature of this conformation. Focused classification revealed positional variations of up to approximately 11 Å, with P domain movements occurring in multiple directions rather than along a single trajectory. Such conformational heterogeneity likely limited the achievable resolution of the P domain region. Notably, the S-domain shell in the rising state was also reconstructed at lower resolution than in the resting state. This may reflect the influence of heterogeneous P domain positioning on particle alignment or the presence of additional structural heterogeneity within the rising state population. Nevertheless, the reconstruction clearly revealed the overall structural rearrangement associated with P domain elevation and rotation away from the shell. Although the map does not permit detailed interpretation of all atomic interactions, it provides important structural insights into the conformational transition and associated reorganization of the P domain interaction network.

Sequence analysis of representative norovirus VP1 proteins revealed varying degrees of conservation among residues involved in the resting- and rising state interaction interfaces (Fig. 6). In the GII.3/TCH04-577 strain analyzed here, the resting site 1 (Re-1), corresponding to the resting state interaction shown in Fig. 4D-1, is located in a relatively exposed region and contains several hydrophilic residues, suggesting a potential role in reversible inter-P domain interactions. The resting site 2 (Re-2) contains the highly conserved DTGR motif and may contribute to stabilization of the resting conformation. The resting site 3 (Re-3) is situated within a proline-rich region containing hydrophobic and aromatic residues, which may impose local structural constraints that help maintain the resting state arrangement. However, these interaction sites are not fully conserved among GI, GII.2, GII.3, and GII.4 noroviruses, indicating that amino acid substitutions may alter the strength or geometry of inter-subunit contacts. In contrast, the Ri site, corresponding to the rising state interaction shown in Fig. 4H, is relatively well conserved among GII.3 and GII.4 strains, particularly with respect to hydrophobic residues. This conservation suggests that the rising site (Ri) contributes to maintaining inter-subunit contacts following P domain elevation and thereby stabilizes the rising conformation. Greater sequence variation in this region among GI.1 and GII.2 strains may therefore result in genotype-dependent differences in the stability of Ri-mediated interactions. Collectively, these sequence variations may influence the conformational equilibrium between resting and rising states, thereby affecting the preferential stabilization of specific conformations in assembled particles. Such differences may also contribute to genotype-dependent variations in recombinant VP1 assembly and the heterogeneous particle populations observed in VLP preparations.

**Figure 6.**
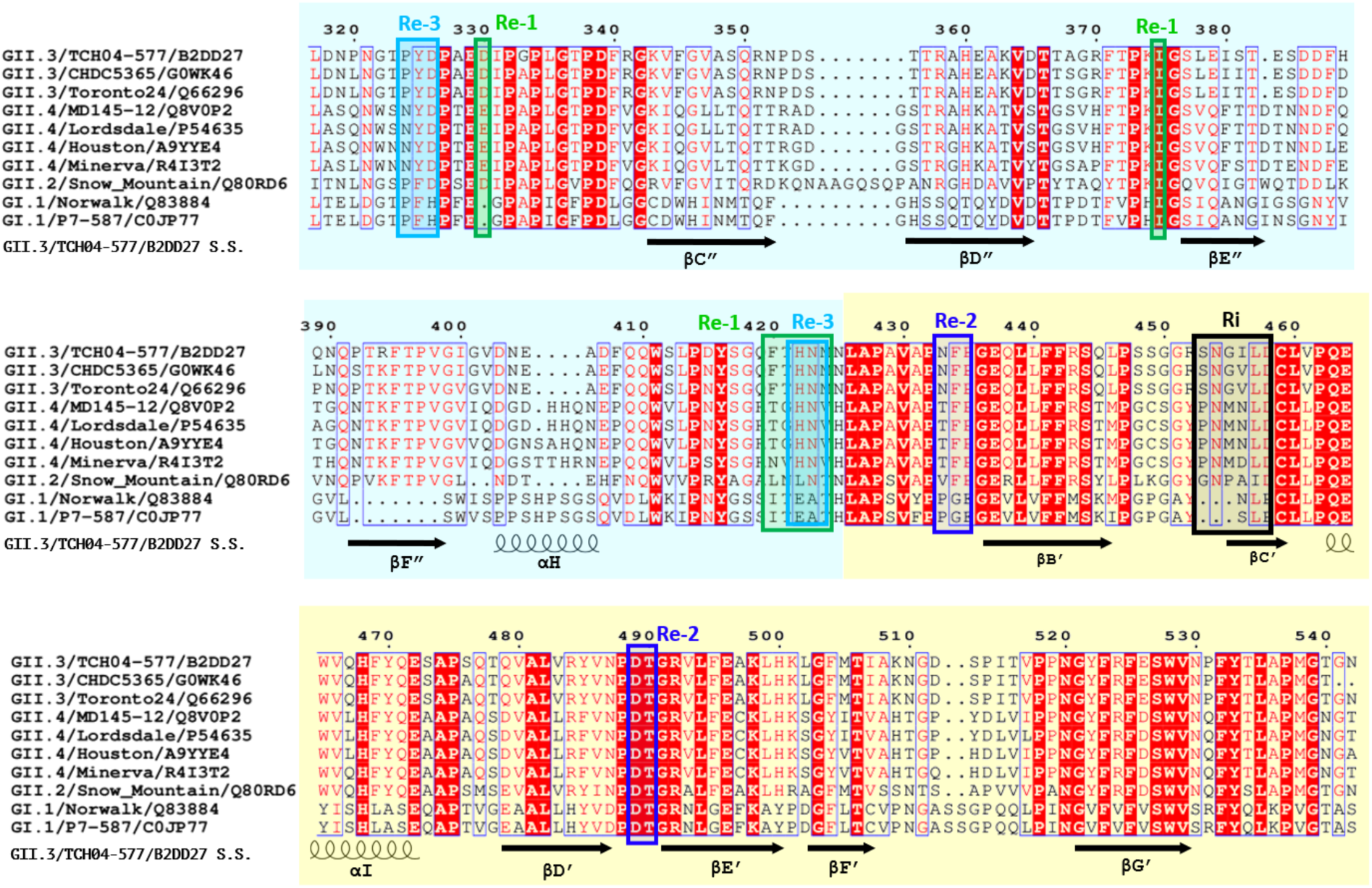
Sequence conservation of VP1 residues involved in resting- and rising state P domain interactions. Multiple sequence alignment of representative norovirus VP1 P domain sequences from GI.1, GII.2, GII.3, and GII.4 strains. The VP1 sequence of the HuNoV GII.3 TCH04-577 strain analyzed in this study is shown at the top. Cyan and pale-yellow shading indicate the P2 and P1 subdomains, respectively. Residues involved in the resting state intermolecular interactions identified in Fig. 4D are designated Re-1 (Fig. 4D-1), Re-2 (Fig. 4D-2), and Re-3 (Fig. 4D-3), whereas residues involved in the rising state interaction identified in Fig. 4H are designated Ri. The Re-1, Re-2, Re-3, and Ri sites are indicated by green, blue, cyan, and black boxes, respectively. The secondary structural elements of the GII.3 TCH04-577 VP1 protein are shown below the alignment. Red shading indicates highly conserved residues, whereas blue outlines denote residues with similar physicochemical properties based on the sequence conservation analysis.

These genotype- and strain-dependent differences in the interaction network may have important functional consequences. The P2 subdomain contains HBGA-binding regions as well as major antigenic sites and receptor-interacting surfaces. Consequently, changes in the position and interaction mode of the P2 region are likely to influence the accessibility of receptor-binding sites and antibody epitopes. In the resting state, the P2-centered interaction network may stabilize an ordered capsid surface optimized for receptor engagement. In contrast, elevation of the P domains in the rising state alters the spacing, orientation, and exposure of the P2 surface while maintaining capsid integrity through P1-mediated contacts. Recent work on GII.4 norovirus entry has suggested that, in addition to HBGA binding, capsid clustering dynamics contribute to functional diversity during the entry process [15]. From this perspective, the transition from a P2-centered resting interface to a P1-centered rising interface may represent a structural mechanism that regulates receptor engagement and subsequent entry-related events.

Furthermore, the observation that broadly neutralizing nanobodies can inhibit GII.4 variants by modulating capsid plasticity [11] suggests that conformational dynamics themselves may constitute an important target for immune recognition and neutralization. The P domain rearrangement identified in HuNoV GII.3 therefore not only advances our understanding of norovirus capsid dynamics but also provides a structural framework for the development of antibody- and nanobody-based strategies aimed at modulating capsid conformational states.

## 4. Materials and Methods

### 4.1 Preparation of HuNoV GII.3 VLP

HuNoV GII.3 TCH04-577 VLPs were produced and purified, as previously described [16,17]. Briefly, the ORF2 region of the HuNoV GII.3 TCH04-577 strain [18] (GenBank accession no. AB365435) was amplified by PCR, cloned into the pDONR221 vector, and subsequently transferred into the pDEST8 vector using the Gateway cloning system (Invitrogen, Carlsbad, CA, USA). Recombinant baculovirus was generated from the resultant plasmid using the BAC-to-BAC expression system (Invitrogen) according to the manufacturer’s protocol.

Recombinant VP1 capsid protein was expressed in High Five insect cells (Invitrogen). VLPs secreted into the culture medium were harvested by centrifugation at 10,000 × g for 30 min and subsequently concentrated by ultracentrifugation at 100,000 × g using an SW32 rotor (Beckman, Palo Alto, CA, USA). Native virion-sized VLPs (approximately 38 nm in diameter) were further purified by CsCl density-gradient ultracentrifugation. The concentration of purified VLP protein was determined using a standard curve generated from serial dilutions of bovine serum albumin (BSA; 0.625-10 μg/μL). For experimental use, VLP preparations were adjusted to a final concentration of 100 ng protein/μL in HEPES buffer. Based on the assumption that each particle contains 180 copies of the 56.6-kDa VP1 capsid protein, 1 ng of VLP protein corresponds to approximately 5.9 × 10^7^ particles. The particle concentration was calculated accordingly.

### 4.2 Cryo-EM data collection for HuNoV GII.3 VLPs

Aliquots (3.0 μL) of the purified HuNoV GII.3 TCH04-577 VLPs were applied to glow-discharged R 1.2/1.3 Quantifoil grids (Quantifoil Micro Tools) coated with a thin carbon support film. Glow discharging was performed immediately prior to sample application using a plasma ion bombarder (PIB-10, Vacuum Device Inc.). The grids were then blotted and plunge-frozen in liquid ethane using a Vitrobot Mark IV (FEI Company) operated at 95% relative humidity and 4°C. Cryo-EM data were collected on a 300-kV Titan Krios transmission electron microscope (Thermo Fisher Scientific) equipped with a Gatan K3 direct electron detector. Images were recorded at a nominal magnification of 53,000×, corresponding to a calibrated pixel size of 1.35 Å at the specimen level. Data were acquired using a low-dose imaging scheme with a total electron exposure of approximately 50 e⁻/Å² distributed over a 5.6 s exposure time. A GIF-Quantum energy filter (Gatan Inc.) with a slit width of 20 eV was used to remove inelastically scattered electrons. Automated data collection was performed using SerialEM software [19].

Detailed data acquisition parameters are summarized in Table S1.

### 4.3 Image processing of HuNoV GII.3 VLPs

All image processing was performed using RELION 4.0 [20]. Movie frames were corrected for beam-induced motion and drift using MotionCor2 [21]. The contrast transfer function (CTF) parameters for each micrograph were estimated with CTFFIND4 (v4.1.5) [22]. Particles were automatically picked from a total of 14,266 micrographs and subjected to reference-free 2D classification (Fig. S1). After the removal of poorly defined classes, 101,338 particles corresponding to T = 3 VLPs were retained for further analysis. Subsequent 2D classification separated the T = 3 particles into two distinct populations corresponding to the resting and rising P domain conformational states. Particles belonging to each conformational state were processed independently through 3D classification and 3D auto-refinement. The final reconstructions were generated from 8,498 particles for the T = 3 resting state and 18,996 particles for the T = 3 rising state. Overall map resolutions were estimated using the gold-standard Fourier shell correlation (FSC) criterion at an FSC threshold of 0.143, based on independently refined half-maps. The final resolutions were 2.8 Å for the T = 3 resting state reconstruction and 7.2 Å for the T = 3 rising state reconstruction.

### 4.4 Atomic model building of HuNoV GII.3 VP1

Cryo-EM maps corresponding to the A, B and C subunits were extracted using UCSF Chimera [23]. An initial atomic model of VP1 was generated by homology modeling based on a previously reported norovirus VP1 structure. The model was manually adjusted to fit the cryo-EM map using COOT [24] and subsequently refined with PHENIX [25].

### 4.5 Focused map refinement of the resting and rising P domains

Focused refinement of the P domains was performed for both the A/B and C/C dimers in the resting and rising VLPs. Symmetry expansion was first carried out using the *relion_particle_symmetry_expand* command in RELION with I1 symmetry applied to the refined particle set. P domain subparticles were then extracted from the symmetry-expanded particles using the map eraser function in UCSF Chimera [23] and the *relion_image_handler* utility in RELION. This procedure generated 150,531 subparticle images for the A/B dimer and 62,943 subparticle images for the C/C dimer in the resting state, and 432,078 subparticle images for the C/C dimer in the rising state. Following 2D classification, the extracted subparticles were subjected to 3D classification without image alignment. Classes exhibiting similar orientations and spatial positions, or a single representative class when appropriate, were selected for further processing and refined using 3D auto-refinement. The focused reconstructions of the resting state VLP yielded resolutions of 3.4 Å for the A/B dimer P domain and 3.1 Å for the C/C dimer P domain. For the rising state VLP, the focused reconstruction of the P domain reached a resolution of 4.7 Å.

## 5. Conclusions

This study demonstrates that the HuNoV GII.3 T = 3 capsid adopts two distinct P domain conformational states, referred to as the resting and rising states. The resting state is characterized by a compact arrangement of the P domains positioned close to the capsid shell surface and stabilized by a P2-mediated interaction network. In contrast, the rising state is characterized by elevation and rotation of the P domains, accompanied by a shift from P2-mediated to P1-mediated inter-subunit interactions. These findings indicate that HuNoV GII.3 capsids possess inherent conformational plasticity and suggest that the equilibrium between resting and rising states may be modulated by genotype- or strain-specific differences in P domain interactions. Such structural plasticity may underlie mechanisms by which HuNoV capsids regulate receptor engagement, evade immune recognition, and adapt to the intestinal environment.

## Author Contributions

Conceptualization, Ka.M.; methodology, M.M., R.T., Ko.M.; validation, C.S.; formal analysis, C.S.; investigation, C.S; resources, M.M and R.T.; data curation, C.S.; writing—original draft prepara-tion, C.S.; writing—review and editing, Ka.M.; visualization, C.S; supervision, Ka.M. and K.K; project administration, Ka.M. and K.K; funding acquisition, Ka.M and K.K. All authors have read and agreed to the published version of the manuscript. Authorship must be limited to those who have contributed substantially to the work reported.

## Data availability statement

Cryo-EM maps for the resting-state overall map, rising-state overall map, resting-state A/B dimer focused map, resting-state C/C dimer focused map, and rising-state C/C dimer focused map have been deposited in the Electron Microscopy Data Bank under accession numbers EMD-81556, EMD-81567, EMD-81571, EMD-81575, and EMD-81576, respectively. The atomic model has been deposited in the Protein Data Bank under accession number 27ZD.

## Acknowledgements

We thank Dr. Mika Hirose and Dr. Takayuki Kato of Institute for Protein Research, Osaka University for collecting data using cryo-EM.

## Abbreviations

2D: two-dimensional
3D: three-dimensional
cryo-EM: cryo-electron microscopy CTF contrast transfer function
FSC: Fourier shell correlation
GI: genogroup I
GII: genogroup II
HBGA: histo-blood group antigen
HuNoV: human norovirus
MNV: murine norovirus
NoV: norovirus
NTD: N-terminal domain
ORF: open reading frame
P: domain protruding domain
P1: protruding 1 subdomain
P2: protruding 2 subdomain
RNA: ribonucleic acid
RMSD: root-mean-square deviation
S: domain shell domain
SPA: single-particle analysis
T: triangulation number
VLP: virus-like particle
VP1: major capsid protein VP1
VP2: minor structural protein VP2

