## Supplemental Table 1-3 and Figure 1-3 for "Conformational plasticity of the human norovirus GII.3 capsid reveals alternative P domain interaction networks"

* Corresponding authors

**Table S1. Data collection, image processing, and model statistics**

| Data collection | |
|---|---|
| Electron microscope | Titan Krios |
| Detector | Gatan K3 |
| Accelerating voltage | 300 kV |
| Magnification | 53,000 |
| Pixel spacing (Å) | 1.35 |
| Exposure time (s) | 5.6 |
| Electron dose rate | 7.14 $e^{-}/Å^{2}/s$ |
| Total electron dose | 50 $e^{-}/Å^{2}$ |
| Defocus range (µm) | 0.5 - 2.5 |

**Table S2. Image processing**

| Image processing | | | | |
|---|---|---|---|---|
| Particle type | T = 3 (resting) | | T = 3 (rising) | |
| Symmetry imposed | I1 | | | |
| Number of micrographs | 14,266 | | | |
| Initial particle number | 101,338 | | | |
| Final particle number | 8,498 | | 18,996 | |
| B-factor applied (Å$^{2}$) | -116 | | -604 | |
| Global resolution (Å) | 2.8 | | 7.2 | |
| EMDB number | EMD-81556 | | EMD-81567 | |
| Focused refinement | A/B dimer | C/C dimer | A/B dimer | C/C dimer |
| Final particle number | 150,531 | 62,943 | - | 432,078 |
| Resolution (Å) | 3.4 | 3.1 | - | 4.7 |
| EMDB number | EMD-81571 | EMD-81575 | - | EMD-81576 |

**Table S3. Model statistics**

| Model building | |
|---|---|
| Type of particles | T = 3 (resting) |
| Number of residues built | 1541 |
| RMSD (bonds) | 0.006 Å |
| RMSD (angles) | 0.904° |
| Ramachandran favored | 96% |
| Ramachandran allowed | 4% |
| Ramachandran outliers | 0% |
| Rotamer outliers | 0.15% |
| C-beta outliers | 0 |
| Clashscore, all atoms | 5 |
| PDB ID | 27ZD |

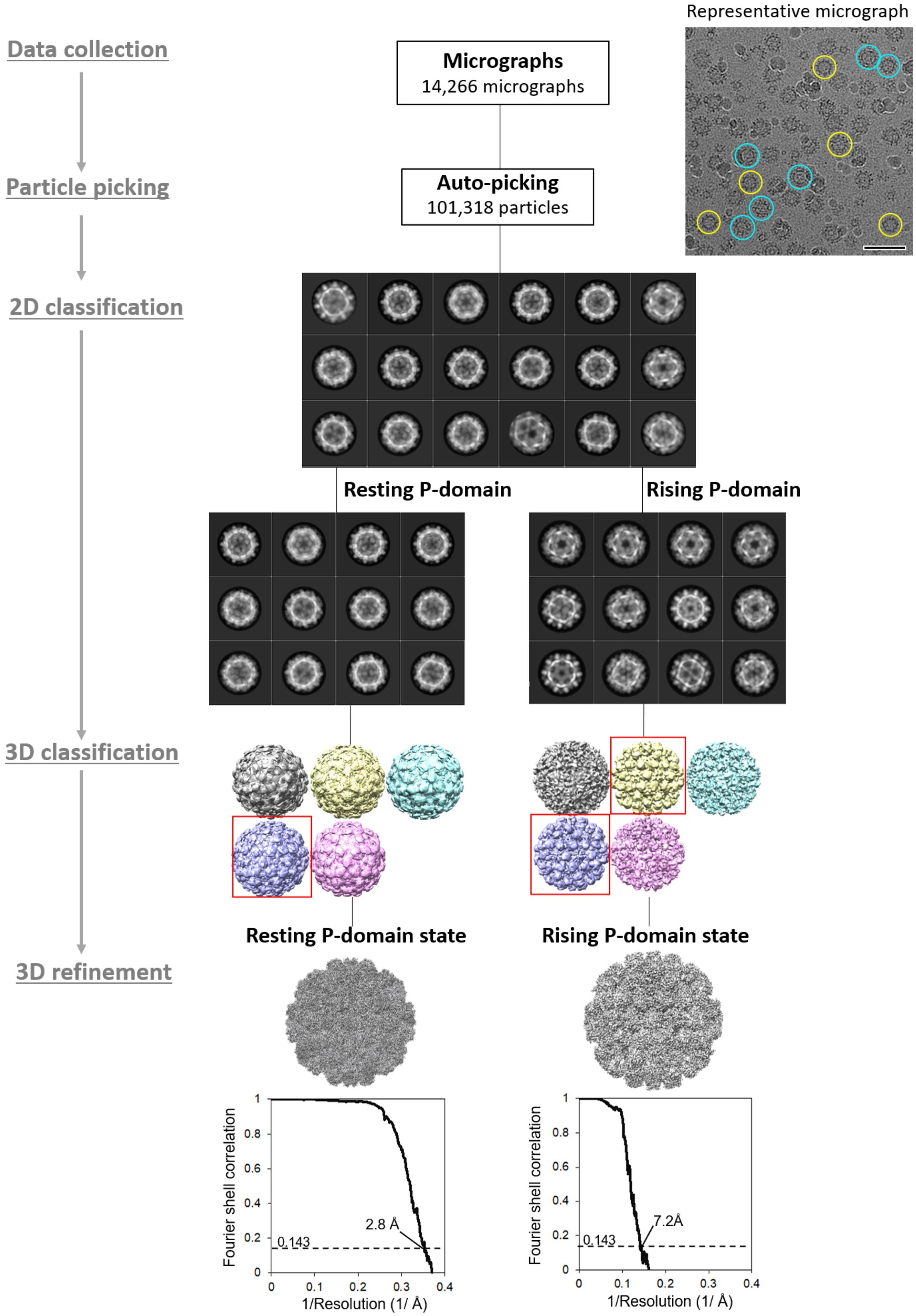
Data collection
Particle picking
2D classification
3D classification
3D refinement
Representative micrograph
Micrographs
14,266 micrographs
Auto-picking
101,318 particles
Resting P-domain
Rising P-domain
Resting P-domain state
Rising P-domain state
Fourier shell correlation
1/Resolution (1/ Å)
0.143
2.8 Å
7.2Å

**Figure S1. Cryo-EM image processing workflow.** The workflow for single-particle cryo-EM analysis of HuNoV GII.3 T = 3 VLPs is summarized. Particles were automatically picked from 14,266 micrographs and subjected to reference-free 2D classification. A total of 101,338 particles corresponding to T = 3 VLPs were refined and further processed through additional classification steps to remove low-quality particles and select homogeneous particle populations. The T = 3 particles were subsequently separated into resting- and rising state populations through an additional round of classification. Final 3D reconstructions were generated from 8,498 particles for the resting state and 18,996 particles for the rising state. Overall map resolutions were estimated using the gold-standard FSC criterion at an FSC threshold of 0.143, yielding resolutions of 2.8 Å for the resting state reconstruction and 7.2 Å for the rising state reconstruction. Corresponding FSC curves are shown at the bottom.

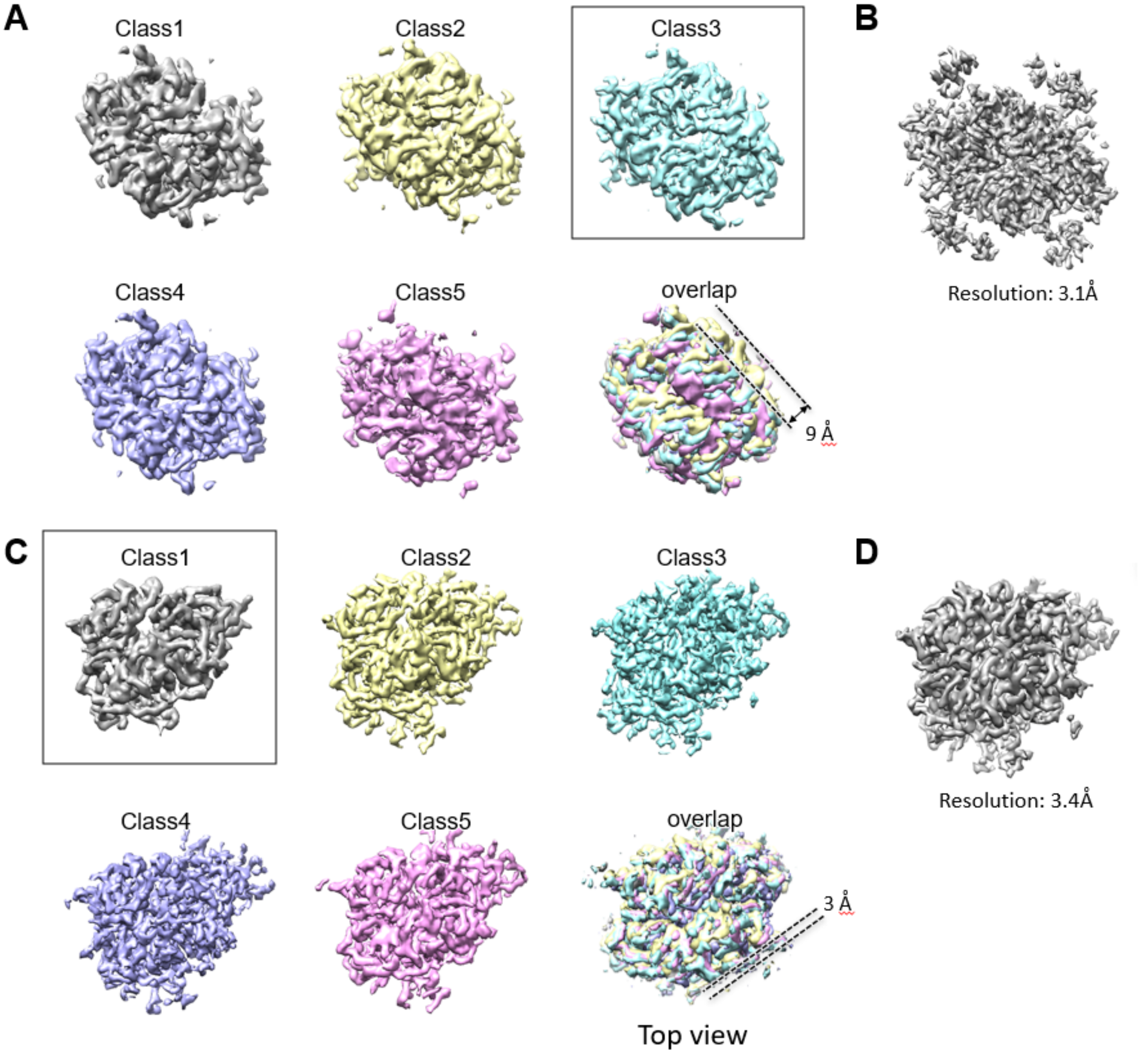


**Figure S2. Focused refinement of the resting P domain dimers in the T = 3 particle.** (A) Representative 3D classes of the resting state P domain in the C/C dimer obtained from symmetry-expanded particles by focused 3D classification without alignment. Because no alignment was performed during classification, each class retains the original spatial position of the corresponding C/C dimer within the T = 3 particle. The class selected for subsequent focused refinement is indicated by a box. The merged view illustrates positional variability of up to approximately 9 Å among the classified C/C dimer maps. (B) Focused-refinement reconstruction of the resting state P domain in the C/C dimer. The number of particles used for reconstruction and the final resolution are indicated. (C) Representative 3D classes of the resting state P domain in the A/B dimer obtained from symmetry-expanded particles by focused 3D classification without image alignment. As no alignment was performed during classification, each class retains the original spatial position of the corresponding A/B dimer within the T = 3 particle. The class selected for subsequent focused refinement is indicated by a box. The merged view illustrates positional variability of up to approximately 3 Å among the classified A/B dimer maps. (D) Focused-refinement reconstruction of the resting state P domain in the A/B dimer. The number of particles used for reconstruction and the final resolution are indicated.

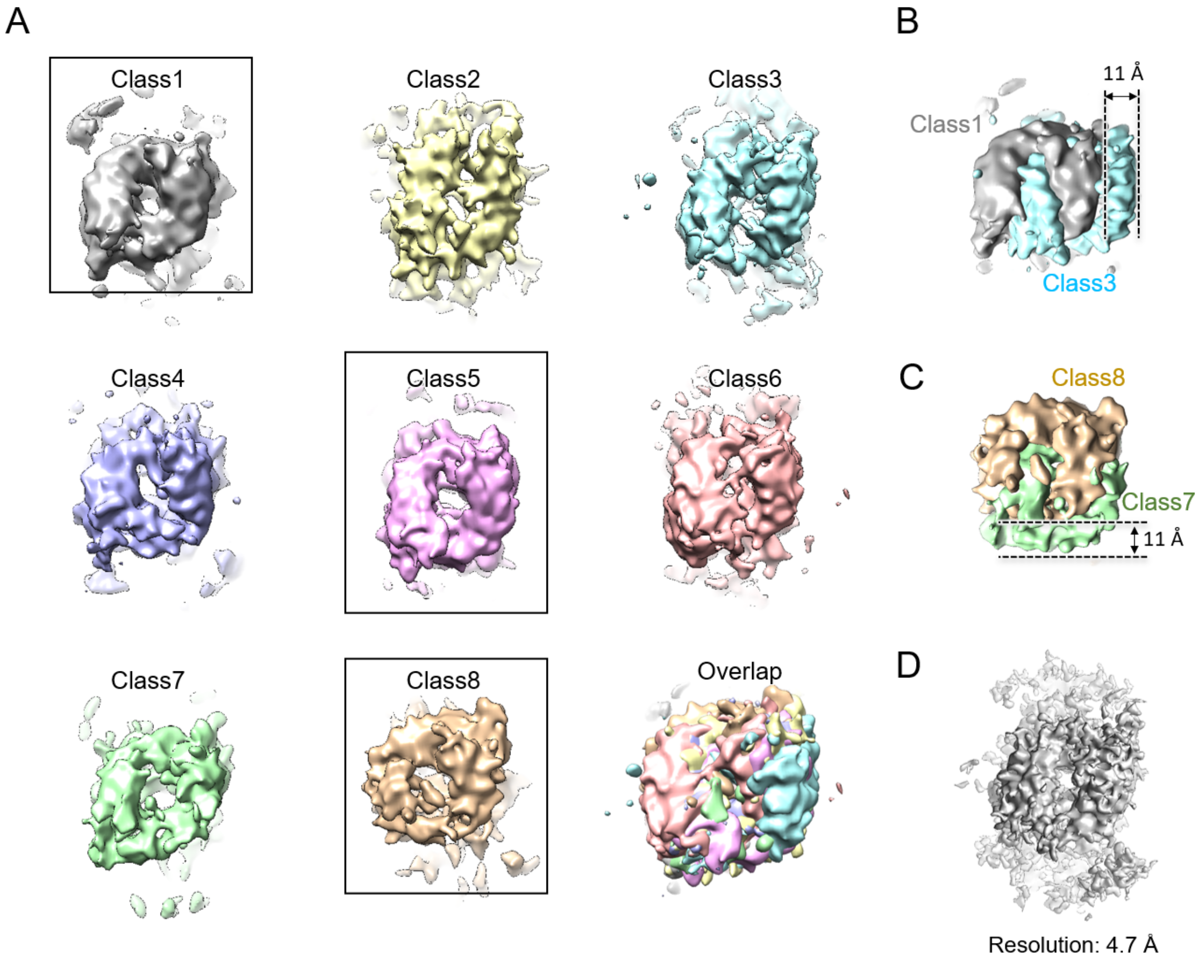


**Figure S3. Focused refinement of the rising P domain C/C dimer in the T = 3 particle.** (A) Representative 3D classes of the rising P domain in the C/C dimer obtained from symmetry-expanded particles by focused 3D classification without alignment. Because no alignment was performed during classification, each class retains the spatial position of the corresponding C/C dimer within the T = 3 particle. Selected classes used for comparison are boxed. The merged view shows positional variability among the classified C/C dimer maps. (B, C) Superimposed views of selected 3D classes showing positional variability of up to approximately 11 Å in different directions. (D) Focused-refinement map of the rising P domain in the C/C dimer. The number of particles and resolution are indicated in the panel.
